# Post-pubertal developmental trajectories of laryngeal shape and size in humans

**DOI:** 10.1101/2022.10.11.511801

**Authors:** Tobias Riede, Amy Stein, Karen L. Baab, Joseph M. Hoxworth

## Abstract

Laryngeal morphotypes have been hypothesized related to both phonation and to laryngeal pathologies. Morphotypes have not been validated or demonstrated quantitatively and sources of shape and size variation are incompletely understood but could be critical for the explanation of behavioral changes (e.g., changes of physical properties of a voice) and for therapeutic approaches to the larynx. Therefore results are likely to have implications for surgeons and speech language pathologists. A stratified human sample was interrogated for phenotypic variation of the vocal organ. First, computed tomography image stacks were used to generate three-dimensional reconstructions of the thyroid cartilage. Then cartilage shapes were quantified using multivariate statistical analysis of high dimensional shape data from margins and surfaces of the thyroid cartilage. The effects of sex, age, body mass index (BMI) and body height on size and shape differences were analyzed. We found that sex, age, BMI and the age-sex interaction showed significant effects on the mixed sex sample. Among males, only age showed a strong effect. The thyroid cartilage increased in overall size, and the angulation between left and right lamina decreased in older males. Age, BMI and the age-height interaction were statistically significant factors within females. The angulation between left and right lamina increased in older females and was smaller in females with greater BMI. A cluster analysis confirmed the strong age effect on larynx shape in males and a complex interaction between the age, BMI and height variables in the female sample. The investigation demonstrated that age and BMI, two risk factors in a range of clinical conditions, are associated with shape and size variation of the human larynx. The effects influence shape differently in female and male larynges. The male-female shape dichotomy is partly size-dependent but predominantly size-independent.

## Introduction

The larynx is a valve located at the crossroads of the alimentary and respiratory tract and is involved in swallowing, breathing and voice production (1,2). All three functions depend on movements that adjust the glottal opening and vocal fold tension. The shape of laryngeal cartilages is one important character that determines how the glottis opens or closes and tension is placed on the vocal folds (3–5). Shape variation of the human larynx is substantial (6–11). The most important source of shape variation is sex (12–14), but some morphometric measures of the larynx are more variable than others even among sexes. An awareness of the anatomical variations of the larynx and their sources is important for surgical approaches to correct glottal insufficiency (15, 16), may inform laryngeal massage therapy (17, 18), and likely helps better understand comparative functional morphology of the larynx in other nonhuman mammals (19, 20). The current study was designed to investigate sources of human laryngeal shape variation. The existence of morphotypes among females or among males was hypothesized in a stratified and balanced human sample. We used multivariate statistics to analyze high-dimensional shape data in the form of landmarks and semilandmarks.

Laryngeal morphometry faces multiple challenges. The complexity of the larynx and the presence of only a few well-defined anatomical landmarks have limited the ability to fully capture small nuances in shape differences. The same reasons have hindered the translatability of findings across species boundaries, for example from animal models to human. Furthermore, measurements taken from cadaver specimens may be inaccurate due to shape changes caused by the post mortem alteration of the tissue or by the excision of the organ from the constraints in the neck. A better understanding of developmental changes in shape and size depends on a sufficiently robust and sensitive approach and the control of confounding variables. Geometric morphometrics is a toolkit of methods for analyzing high dimensional shape data in the form of surface landmarks and semilandmarks capable of identifying even subtle differences in form (21). The approach used here relies on a mix of homologous, anatomical landmarks and surface semilandmarks whose positions are loosely anchored by these few fixed landmarks, all taken on 3D surface reconstructions of laryngeal cartilages. Moreover, the sample consists of clinical computed tomography scans from living patients, avoiding issues specific to cadavers, and, for the first time, providing sufficiently large samples to investigate the role of aging and body composition on larynx anatomy.

### Background

Sex is the biggest source for shape and size variation of the human larynx (7–10, 12–14, 22, 23) and has consequences for its function (24–26). Laryngeal measures differ from 10 to 50% between adult male and females (7–10, 27). This sexual dimorphism results from an individual’s endocrine profile during puberty (28–33). What has remained elusive is the relative contribution of other genetic and environmental factors after puberty. The post-pubertal development of larynx shape and size between ages 18 and 60 has received little attention. Previous investigations have focused on populations of advanced age (8) because characteristic voice problems of the elderly population often set in after the age of 60 (34, 35). Factors that contribute to vocal changes at advanced age include structural changes of the vocal fold, innervation deficits of intralaryngeal muscles and structural changes of the cartilaginous framework (36).

A problem for laryngeal morphometry is the use of linear measurements, many of which are not ideal for biomechanical modeling purposes (27). Computational modeling of laryngeal movements (4, 24) represents an important tool for the translational process (i.e., the development of possible cures for diseases) and for the evaluation of the relationship between form and function of the larynx (37). Finite element models of larynx and vocal fold tissue can provide a functional explanation of shape differences which is an important prerequisite for a translation of morphometric results to clinical relevance (38, 39). The current study represents an improvement in how we quantify morphology of the laryngeal framework and a more complete characterization of biological influences on the larynx. The resulting surface renderings can be interpreted with respect to phonation, bridging the current gap between phenotypic shape and function.

Work in mice demonstrated that the laryngeal cartilages change in size and shape throughout life and the four main laryngeal cartilages do not develop uniformly (40). The purpose of the current study was to investigate whether age and body mass, two important risk factors in many clinical conditions, contribute to laryngeal shape changes in women or in men throughout life. The results of this study have important implications for surgeries of the larynx. Surgical manipulation of the cartilaginous framework such as thyroplasty (15, 16, 41, 42) or reconstructive procedures of the cartilage framework (42–46) are important treatment options to re-establish glottal sufficiency. Identifying the relative contributions of sex, age and body mass index (BMI) to shape variation of the laryngeal cartilaginous framework will be informative for surgical planning.

Finally, we expect to learn more about developmental trajectories of larynx form and function. For example, the existence of laryngeal morphotypes (i.e. a group of individuals within a sex with similar morphologies) has been speculated (6, 47), and may predispose people to certain pathologies. If we can identify morphotypes, we expect that those can be linked to certain sexes, ages and possibly other genetic or environmental factors.

## Results

### a) Shape variation

The first two shape axes of the mixed sex sample accounted for 31.7% and 10.9% of the explained variation, respectively (Figure 1A) and regressions of PC scores on PC 1 and 2 on sex are significant (Table 2). Variation among individuals along the first principal component (PC1) mostly reflects male-female differences (Figure 1A). While the first PC reflects size differences (regression of PC 1 scores on centroid size: R^2^= 0.40, p<0.001), this is driven by the fact that males are larger than females on average; the regressions of PC scores on cartilage size within each of the sexes are statistically significant for females (R^2^= 0.047, p=0.026) but not for males (R^2^= 0.009, p=0.33) and do not explain much of the variation even in females. In other words, individuals with a larger thyroid cartilage do not differ from individuals of the same sex with a smaller cartilage in the same direction that males differ from females. The primary axis of shape variation in the thyroid cartilage relates to the angulation of the right and left thyroid laminae, the orientation of the superior border of the thyroid cartilage and the relative height of both laminae (Figure 1B). Lower scoring individuals, which are predominantly female, have a thyroid cartilage that is anterior-posterior (AP) compressed and relatively wide superiorly with a superior border that is angled upward ventrally (Figure 1B). The second PC of thyroid cartilage shape contrasts individuals with an AP compressed cartilage and a long superior horn and those with a more AP expanded cartilage and a shorter superior horn (Figure 1C). Unlike variation along PC 1, variation along PC 2 does not correspond to male-female differences (Figure 1A).

**Figure 1:**
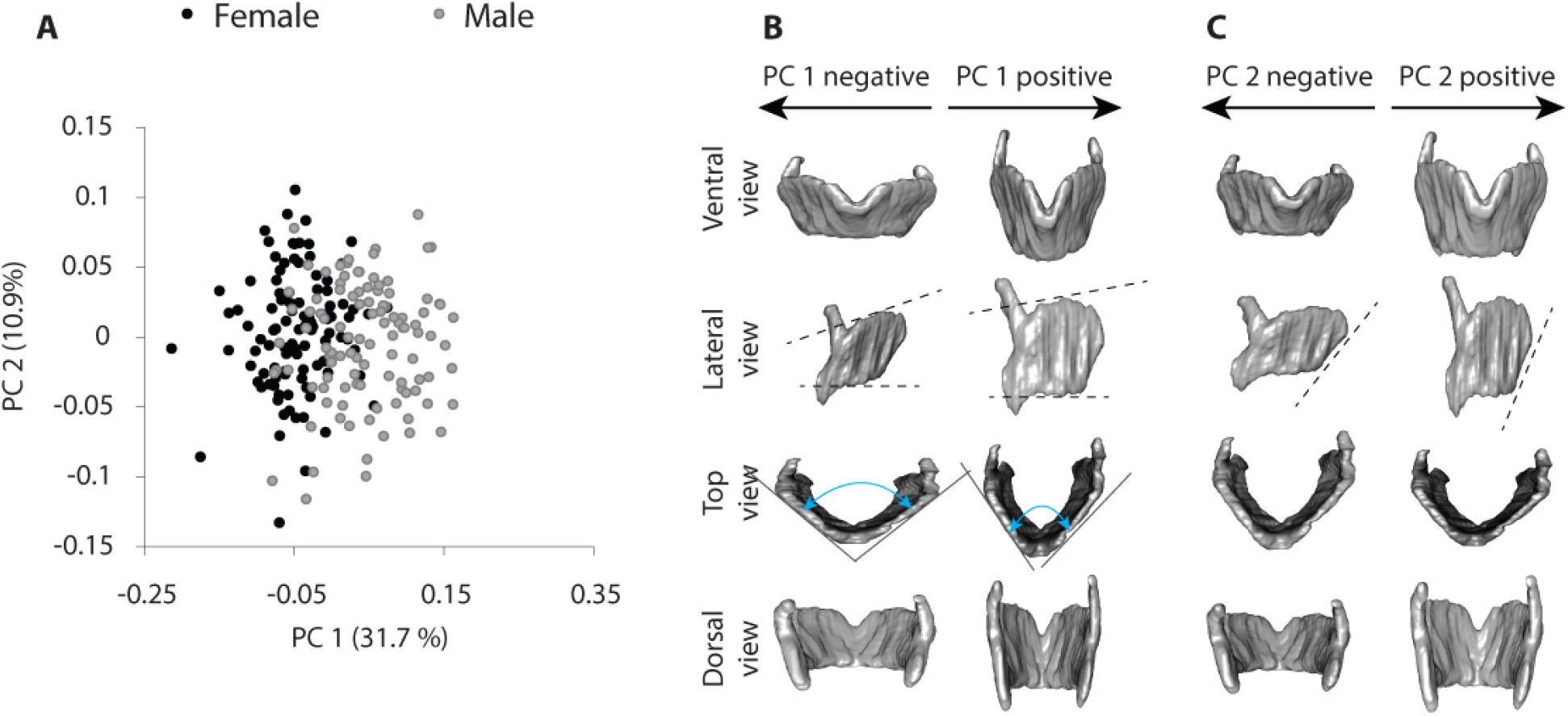
Principal component analysis of shape coordinates (PC1 and PC2) from landmarks. Each subjects takes a specific point in the two-dimensional space of PC1 and PC2 (**A**). Warped cartilage images showing shape differences in the thyroid cartilage for the first (**B**) and second shape axis (**C)**. Images represent shape differences associated with the mean minimum and maximum extents of PC 1 and PC 2. **C**: Ordination of first 2 PC scores of Procrustes shape coordinates for the thyroid cartilage. Note a considerable overlap between the sexes on the first shape axis and a complete overlap on the second.

### b) The correlation among sex, age, BMI, body height and cartilage size

Three independent variables, age, sex and BMI, were uncorrelated with each other (Figure 2 A-C). The cartilage was larger on average in males than females (Figure 2 D). There was no correlation between age and cartilage size (Figure 2 E) and no correlation between BMI and cartilage size when both sexes were combined (Figure 2 F). Unexpectedly, centroid size was positively correlated with age in males (Figure 2 E). Average centroid size was about 4% larger in the 78 to 88-year age group than in the 18 to 28-year age group suggesting a growth rate of about 0.67% per decade. Females showed no correlation between centroid size and age (Figure 2 E).

**Figure 2:**
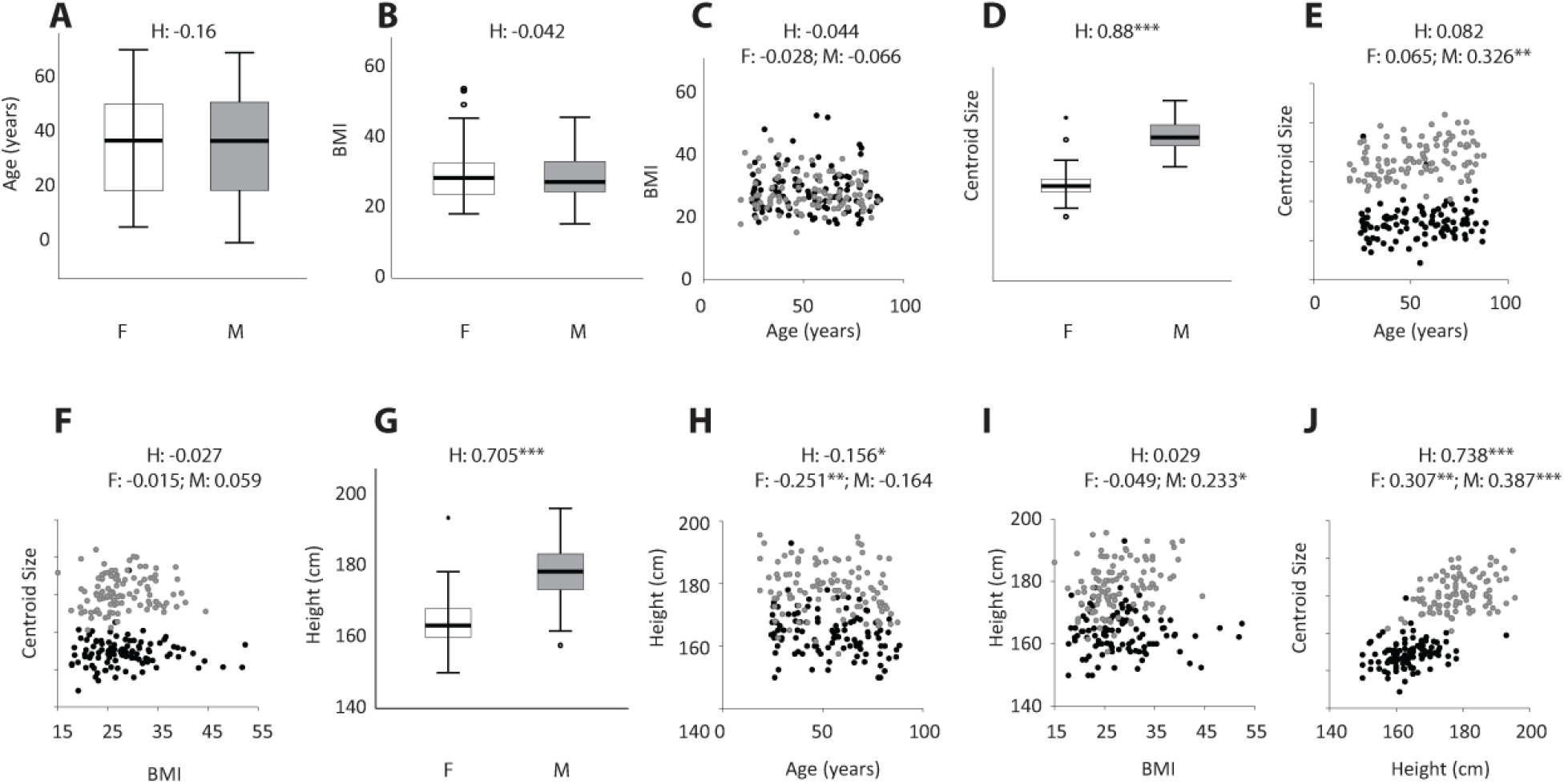
Correlations illustrating the relationships between sex, age, BMI, centroid size and body height. The sample included 105 females and 105 males which were stratified according to age and BMI (**A**, **B**, **C**). The cartilage size (‘centroid size’) was larger in males (**D**) and increased throughout life in males but not in females (**E**). Centroid size and BMI were not associated (F). Males were taller than females (**G**), and older females were shorter on average (**H**). BMI was higher in taller males (**I**). The cartilage size (‘centroid size’) was associated with body height in both sexes (**J**). H: mixed-sex sample; F: female sample; M: male sample; Significance codes: 0.001 ‘***’ 0.01 ‘**’ 0.05 ‘*’

Males were taller than females (Figure 2 G). While body height was weakly associated with age and BMI (Figure 2 H, I), body height, as expected, showed a strong correlation with cartilage size in the mixed sex sample; the relationship between height and cartilage size is stronger in males than females (Figure 2 J). Body height explained neither the increase in larynx size in aging men (partial correlation controlling for height r = 0.428, p < 0.001), nor the absence of such a relationship in females (r = 0.159, p =0.106) as expected given that individuals typically reach their adult height by 18 years of age.

### c) The relationship of larynx shape to sex, age, BMI and centroid size

The full MANOVA model including age, sex and BMI as main effects, and their two-way interactions, accounted for 20.3% of shape variation in thyroid cartilage shape (19.2% when the non-significant interactions were dropped from the model) (Table 1). There was a significant interaction between sex and age, which we explore in greater detail below, but it accounted for only 0.8% of the shape variation. The most significant factor affecting thyroid shape was sex, which accounted for 14.9% of variation independent of the influence of age and BMI.

**Table 1:** Multiple Analysis of Variance of thyroid cartilage shape with significance testing using Residual Randomization method, 1000 permutations, Ordinary Least Squares, Type II model, Effect sizes (Z) based on F distributions. Significance Codes: 0.001 ‘**’ 0.01 ‘*’ 0.05

|  | **Df** | **SS** | **MS** | **R<sup>2</sup>** | **F** | **Z** | **P** |
| --- | --- | --- | --- | --- | --- | --- | --- |
| Sex | 1 | 0.5046 | 0.50462 | 0.1492 | 37.88 | 8.41 | 0.001 ** |
| Age | 1 | 0.0849 | 0.08489 | 0.0251 | 6.37 | 5.40 | 0.001 ** |
| BMI | 1 | 0.0396 | 0.03963 | 0.0117 | 2.97 | 3.25 | 0.005 ** |
| Sex:Age | 1 | 0.0303 | 0.03034 | 0.0090 | 2.28 | 2.45 | 0.014 * |
| Sex:BMI | 1 | 0.0127 | 0.01270 | 0.00375 | 0.95 | 0.02 | 0.475 |
| Age:BMI | 1 | 0.0193 | 0.0193 | 0.00569 | 1.45 | 1.24 | 0.112 |
| Residuals | 203 | 2.7043 | 0.01332 | 0.79944 |  |  |  |
| Total | 209 | 3.3827 |  |  |  |  |  |

**Table 2:** Regression of shape axis on sex. Significance Codes: ‘***’ 0; ‘*’ 0.05

| Shape axis | Unstandardized | Unstandardized | Standardized | t | Sign. |
| --- | --- | --- | --- | --- | --- |
| PC1 | 4.623 | .289 | .661 | **16.017** | **.000\*\*\*** |
| PC2 | -1.110 | .492 | -.093 | **-2.255** | **.025\*** |

### d) Sex specific variation in cartilage shape

Next, we evaluated the shape variation in the thyroid cartilage for each sex. The primary axis relates to the angulation of the right and left thyroid laminae and the orientation of the superior border of the thyroid cartilage within both males and females. The angulation was larger and the superior border points cranially in the lower scoring individuals in both sexes (Figure 3A, B). High scoring individuals in each sex have a thyroid cartilage that is more AP compressed and relatively wider superiorly with a superior border that is angled upward ventrally (Figure 3 A, B). Lower scoring individuals in each sex demonstrate tighter angulation between the left and right thyroid lamina. In males, the lamina height was greater in high scoring individuals. The second PC of thyroid cartilage shape captures variation in the relatively length of the superior horn in both sexes (Figure 3 A, B). In females, the second axis contrasts also differences in relative AP distance (Figure 3A).

**Figure 3:**
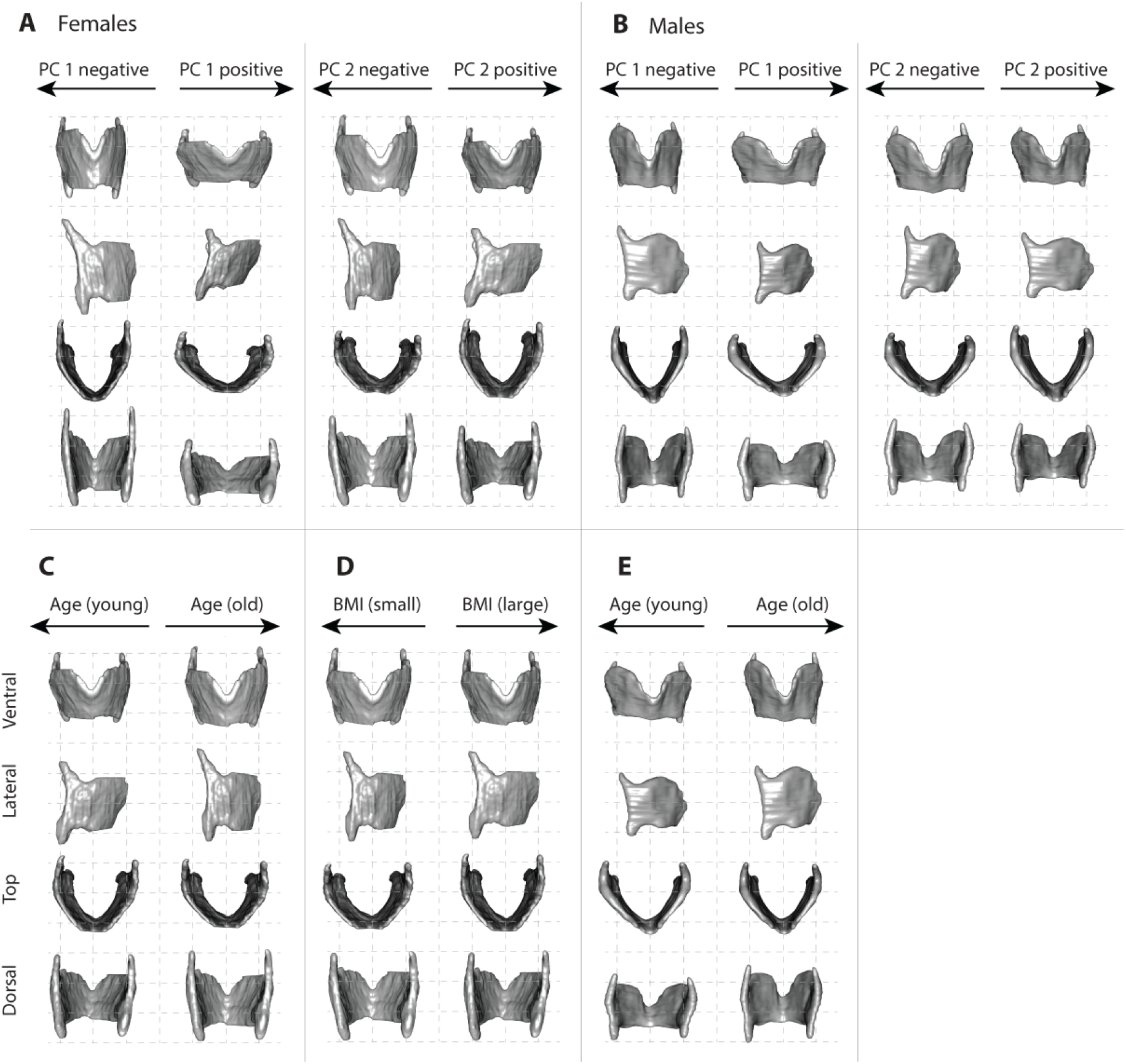
Warped thyroid cartilage images showing shape variation within females (**A**) and within males (**B**) for the first and second shape axis of the single sex PCAs. In both females and males, individuals on the first shape axis are differentiated by the angulation between the left and right lamina. Images in **C** through **J** illustrate thin-plate-spline warps of regression models for age (**C**, **E**) and BMI (**D**, **F**) on a mean female and male surface rendering, respectively. The procedure first selects the individual that is closest to the group (e.g., female, male) average. Then the group average was warped toward the two ends of the slope (regression line) defined by the regression coefficients of the two models. Shape differences were exaggerated by a factor of 2 to make them more visually prominent. Note that age and BMI were significant factors for females but only the model for age was significant for males (Table 3).

Next, we evaluated the effect of age, BMI, centroid size and height on cartilage shape within each sex separately. We used the same MANOVA model described above but removed the sex variable and added the height variable, with pairwise interactions again included in the model. The full model accounted for 11.2% and 8.4% of shape variation in the female- and male-only analyses, respectively (Table 3). The interaction between age and height, and the main effects of age and BMI were significant in females accounting for 1.9%, 4.2% and 1.9%, respectively (Table 3). In males, only age was a significant factor, explaining 3.3% of shape differences.

**Table 3:**
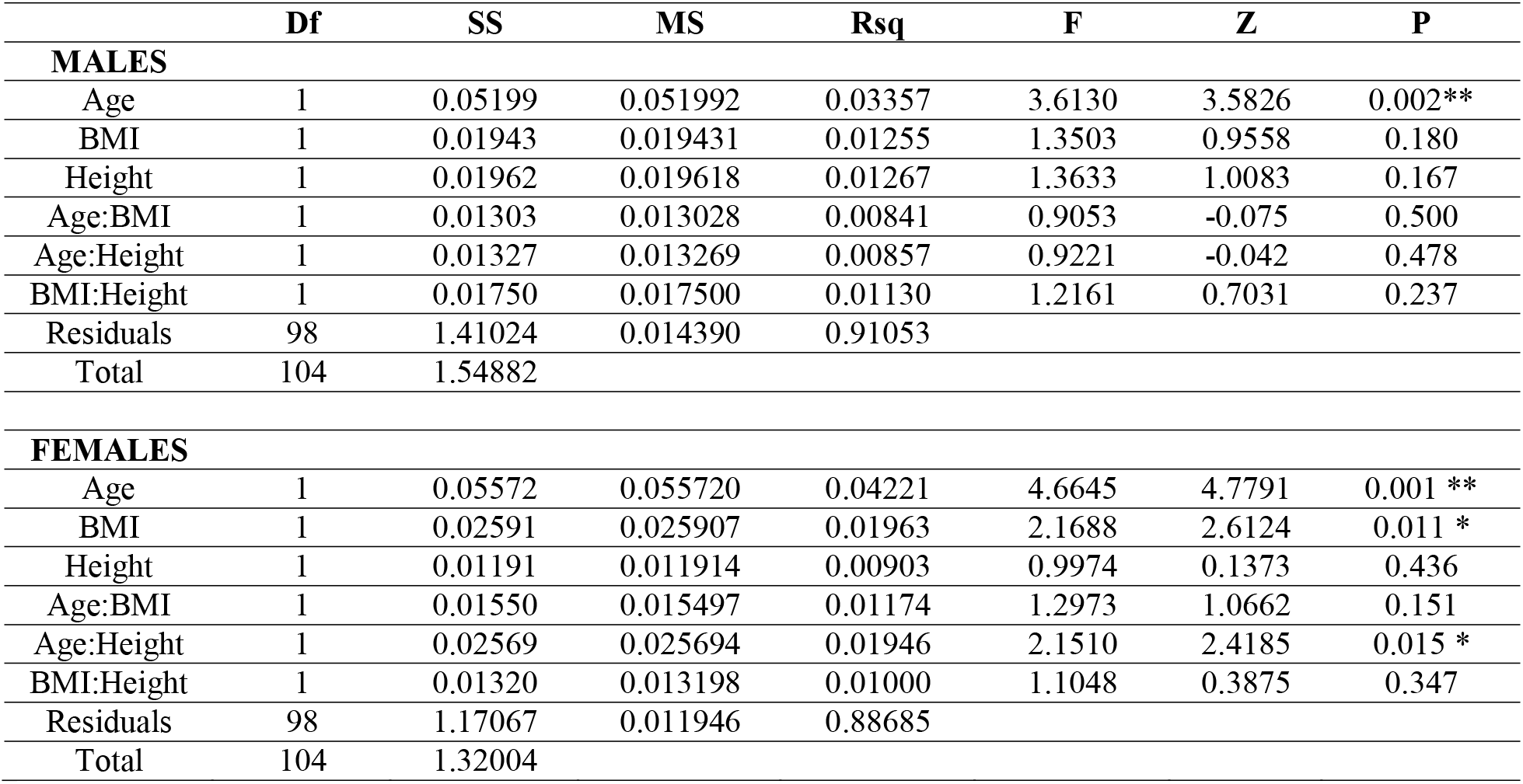
Multiple Analysis of Variance of thyroid cartilage shape separate for males and females and using a Residual Randomization method, 1000 permutations, Ordinary Least Squares, Type II, Effect sizes (Z) based on F distributions. Signif. Codes: 0.001 ‘**’ 0.01 ‘*’

We illustrated shape variation associated with increasing age for each sex in Figure 3 C-F since the interaction between sex and age was significant in the mixed-sex model, and age played a role in both sex-specific models. In older females, the angulation between left and right thyroid lamina increases (i.e. thyroid cartilage is AP compressed) and the superior border becomes angled upward ventrally (Figure 3C). The thyroid cartilage is also more AP compressed in females with a smaller BMI, i.e. it appears as if older age and larger BMI have opposite effects on the angulation in females (Figure 3D). We found that the angulation between left and right thyroid lamina decreases in older males (Figure 3E) and the laminae are relatively more expanded cranio-caudally (Figure 3E). The effect of BMI appears to be similar in males and females, however note that BMI only reached statistical significance in females (Table 3).

In order to translate statistical findings into functionally relevant information, we investigated five measures that were previously identified as biomechanically relevant (Figure 4A). Those five measure were taken from the warped extremes for age, BMI, centroid size and height. Note that only age was significant for males and only age and BMI were significant for females (Table 3). Figure 4B illustrates direction and magnitude of the four effects on each measure.

**Figure 4:**
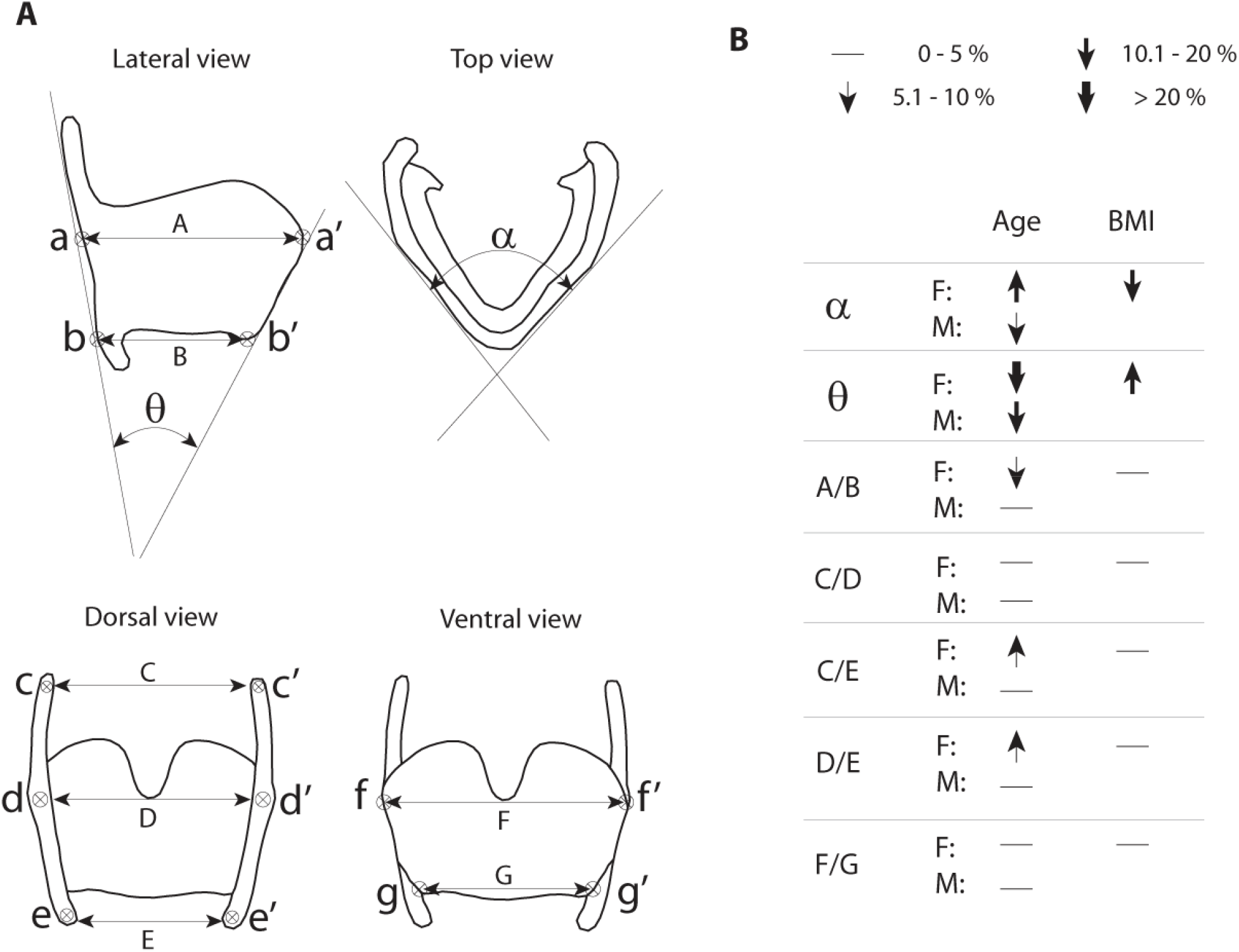
Translating shape variation into biomechanically relevant information. (**A**) Numerous measures have been previously identified as biomechanically relevant for laryngeal movements (Tayama et al. 2001). Seven linear measures and two angles were measured in the current data set using regression models for age and BMI (illustrated in Figure 3). (**B**) Arrow direction indicates the direction of change of the measure with an increase in age and BMI, respectively. Arrow thickness indicates the magnitude of change between the 18-to-28-years age group and the 78-to-88-years age group.

### f) Cluster analysis

Next, we explored different morphotypes first within the mixed-sex sample and then among females and males. Using k-means cluster analysis, the 2-cluster solution for the mixed-sex sample separated most males (75.3%) into Cluster 1 and most females (87.1%) into Cluster 2 (k-means clustering; silhouette coefficient = 0.23; 20 shape axes) (Figure 5A). Individuals in Cluster 1 were taller than in Cluster 2, and their larynx size was larger (consistent with typical sex differences) (Table 4). There were no differences in age and BMI between the two clusters (Table 4).

**Figure 5:**
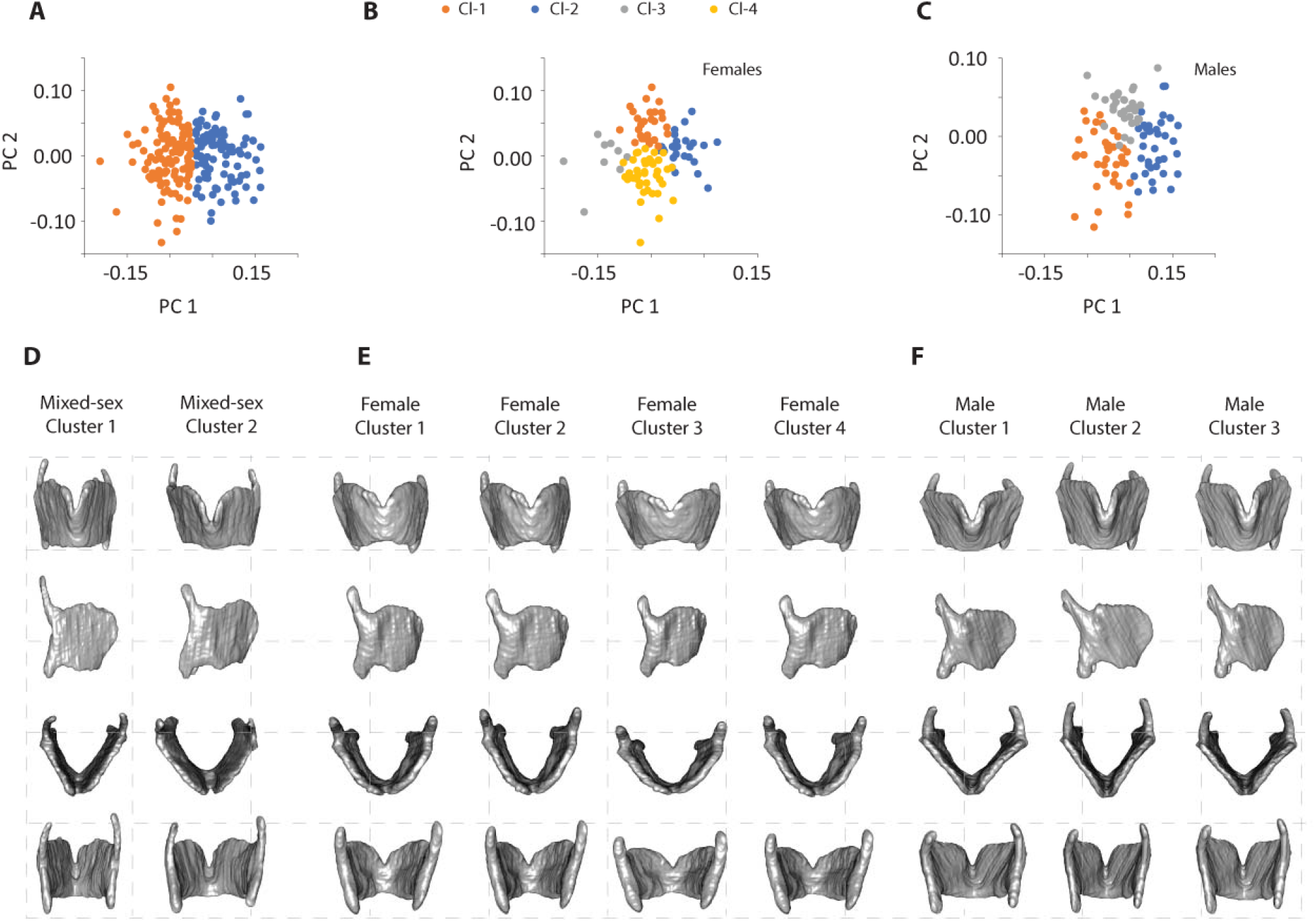
Cluster analysis for mixed-sex (**A**), female (**B**) and male (**C**) data sets. Images in **D** through **F** illustrate thin-plate-spline method warps of regression models for cluster averages. The respective population average (mixed-sex, male or female) was warped to the shape of each cluster average.

**Table 4:** Comparison of mean (standard deviation) values for the age, BMI, centroid size and body height variables for individuals classified into Clusters 1 and 2, respectively, based on the mixed sex sample. P values indicate that the means between the two cluster were significantly different for centroid size and body height but not for age and BMI. Signif. Codes: <0.001 ‘***’

|  | **Cluster 1 (n = 91)** | **Cluster 2 (n = 119)** | P-value |
| --- | --- | --- | --- |
| Age | 56.2 (19.3) | 51.55 (18.9) | 0.0796 |
| BMI | 28.66 (5.82) | 27.72 (6.73) | 0.2872 |
| Thyroid Size | 238517 (20006) | 201983 (21017) | <0.001*** |
| Height | 177.23 (9.6) | 166.64 (8.6) | <0.001*** |

Next we investigated characteristics of the males and females in both clusters. Males that clustered with females in Cluster 2 (26 out of 105) were significantly younger, of shorter stature and had smaller thyroid cartilages than that of those males in Cluster 1. Indeed, Cluster 1 could be more accurately described as containing the larger individuals of each sex and Cluster 2 includes smaller individuals of each sex, even though the size difference did not reach signficance within females (Table 5). The difference between the average thyroid cartilage shapes in the two clusters (Figure 5D) resembles the previously described sex differences (Figure 1C). In Cluster 2, individuals have a thyroid cartilage that is AP compressed and relatively wide superiorly with a superior border that is angled upward ventrally (Figure 5D).

**Table 5:** Comparison of mean values (standard deviation) for age, BMI, thyroid CS and body height for males and females classified into Clusters 1 and 2 based on the mixed sex sample. Note that the majority of subjects in Cluster 2 were male and the majority in Cluster 1 were female. The average (standard deviation) of each metric in Cluster 1 and 2 was compared between the incorrectly clustered group for males and females separately. Two-sample t-tests suggest that males which were classified into cluster 2, tended to be significantly younger, smaller and with smaller centroid size. For females, none of the variables showed a difference between Cluster1 and 2 averages.

|  | Males (N=105) |  |  | Females (N=105) |  |  |
| --- | --- | --- | --- | --- | --- | --- |
|  | Cluster 2<br>N=26 | Cluster 1<br>N=79 | P | Cluster 2<br>N=93 | Cluster 1<br>N=12 | P |
| Age | 46.2 (19.3) | 55.62 (19.0) | 0.035* | 53.06 (18.6) | 60.27 (21.7) | 0.2918 |
| BMI | 26.39 (5.2) | 28.36 (5.7) | 0.11 | 28.10 (7.1) | 30.72 (6.5) | 0.2136 |
| Thyroid<br>CS | 235952<br>(12856) | 243816<br>(133322) | 0.010* | 192487<br>(10216) | 203634<br>(22071) | 0.1109 |
| Height | 176.00 (6.9) | 179.44 (8.0) | 0.04* | 162.67 (6.4) | 164.03 (7.1) | 0.5019 |

When the k-clustering analysis was performed on females only, there were four female clusters recovered that differed in age, BMI, and thyroid size (Figure 5B, Table 6). Females in Cluster 4 were 46.4 years old on average, showed the highest BMI but smallest centroid size. Cluster 3 is the smallest cluster with only 9 individuals. This cluster had the highest average age (65.3 years) of the 4 clusters and a low average BMI (together with Cluster 1). Females in Clusters 1 and 2 were intermediate in age (59.5 and 55.6 years on average, respectively) and females in Cluster 1 had a low average BMI. Figure 5E illustrates shape differences between the 4 female clusters. The angulation between left and right lamina is wider, and the degree of AP compression is more prominent in Cluster 3, which contains individuals with the highest average age.

**Table 6:**
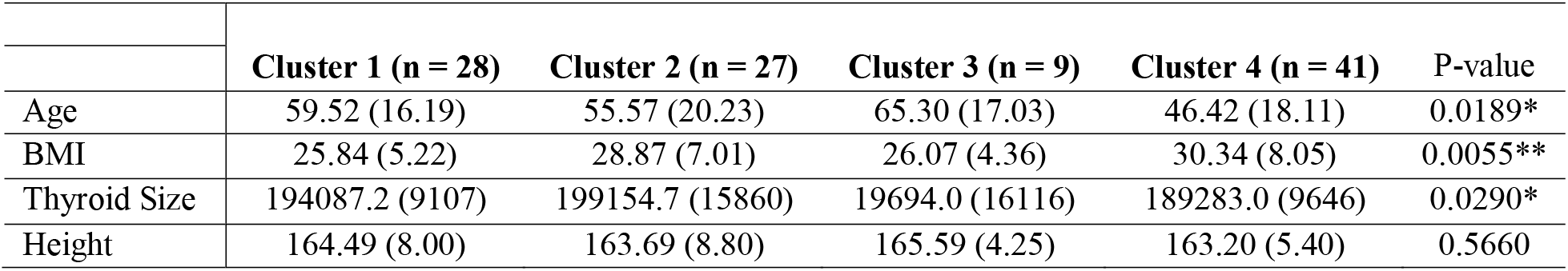
Mean values (standard deviation) for age, BMI, thyroid size and body height variables for females in different clusters. P values indicate that the means between clusters were significantly different for age, BMI and centroid size but not for body height. Signif. Codes: 0.001 ‘**’ 0.01 ‘*’

|  | **Cluster 1 (n = 28)** | **Cluster 2 (n = 27)** | **Cluster 3 (n = 9)** | **Cluster 4 (n = 41)** | **P-value** |
| --- | --- | --- | --- | --- | --- |
| Age | 59.52 (16.19) | 55.57 (20.23) | 65.30 (17.03) | 46.42 (18.11) | 0.0189* |
| BMI | 25.84 (5.22) | 28.87 (7.01) | 26.07 (4.36) | 30.34 (8.05) | 0.0055** |
| Thyroid Size | 194087.2 (9107) | 199154.7 (15860) | 19694.0 (16116) | 189283.0 (9646) | 0.0290* |
| Height | 164.49 (8.00) | 163.69 (8.80) | 165.59 (4.25) | 163.20 (5.40) | 0.5660 |

Three male clusters differed significantly in age and centroid size (Figure 5C, Table 7). The youngest individuals with the smallest larynx were in Cluster 1. Individuals in Cluster 3 were oldest and their centroid size was largest. Age and centroid size of individuals in Cluster 2 were intermediate between Clusters 1 and 3. The angulation between left and right lamina was highest in Cluster 1 and lowest in Cluster 3. Interestingly the oldest/largest and smallest/youngest are not the most different in shape. Figure 5F illustrates shape differences between the 3 male clusters. The most different are the young/small vs. intermediate individuals. The angulation between left and right lamina is wider in clusters 1 and 3. The height of the laminae is taller in Clusters 2 and 3.

**Table 7:** Mean values (standard deviation) for the age, BMI, thyroid size and body height variables for males in different clusters. P values indicate that the means between clusters were significantly different for age and centroid size but not for BMI and body height. Signif. Codes: <0.001 ‘***’

|  | **Cluster 1 (n = 37)** | **Cluster 2 (n = 37)** | **Cluster 3 (n = 31)** | **P-value** |
| --- | --- | --- | --- | --- |
| Age | 43.91 (18.46) | 55.67 (18.49) | 61.60 (17.31) | <0.001*** |
| BMI | 28.00 (6.08) | 28.63 (5.10) | 26.80 (5.66) | 0.4100 |
| Thyroid Size | 236676 (13030) | 241526.5 (10854) | 248475.0 (14692) | <0.001*** |
| Height | 178.72 (8.02) | 178.86 (7.54) | 178.12 (8.20) | 0.7640 |

## Discussion

A strength of this study is the use of a large stratified sample that allows for the statistical separation of multiple variables. The analysis of thyroid cartilage confirmed sex as a major source of size and shape variation. Organ size was significantly different between sexes but did not explain shape variation among the sexes, instead our results support the hypothesis that additional sources of shape variation, for example age and BMI, exist within sexes. Four findings especially inform the understanding of postpubertal shape changes of the human larynx. First, the angulation between left and right lamina is a major shape difference within each sex, but also separates the average male and female shape. This does not appear to be a simple issue of size-shape (i.e., allometric) scaling. Second, the male larynx demonstrates a life-long size increase, on the order of ~0.7% per decade. Third, age has a significant effect on thyroid cartilage shape within males and females, as do BMI and the age-height interaction in females only. Post-pubertal changes in larynx shape differ between males and females and the shape changes associated with sex and aging have biomechanical consequences. Fourth, cluster analyses identified at least four morphotypes among females that correspond to differences in age, BMI and centroid size. In males, cluster analyses identified three morphotypes which differ by age and larynx size.

Next, we discuss the relevance of the four findings for larynx function and hypothesize that life-long post-pubertal trajectories of laryngeal shape changes are causally affected by BMI and body size and are, in part, responsible for functional changes of the larynx.

### Patterns in the mixed-sex sample

The variables examined in the current study (sex, age, BMI and their respective interactions) explain about a fifth of the total shape variation in the thyroid cartilage. Even taking BMI and age into account, 15% of shape variation in the mixed-sex sample is attributable to sex. The BMI, age and height variables account for about 10% of total variation (11.2% and 8.4%, respectively in the female and male samples) in the sex-specific MANOVA models. Other genetic and environmental factors explain the remaining 80% of the shape variation.

The shape differences along the first principal component axis were associated with variation in sex. The shape difference concerns the well-known angulation between left and right lamina and the height of the laminae (12). Males have relatively longer (antero-posteriorly), taller and more acutely angled thyroid cartilages (Figure 3A,B). Interestingly, the primary axis of shape variation within the male and female samples shares similar shape differences to that seen in the mixed-sex sample. On the surface this appears to be a case of size-shape scaling (i.e., allometric shape differences) given that all males are, with few exceptions, larger than any female in our sample. However, this scaling effect does not hold within either sex: individuals with larger cartilages do not score higher on PC 1 in the mixed-sex analysis than those with smaller cartilages within either sex. This hints at factors other than simple scaling driving the sex-differences.

To further disentangle levels of variation, future study designs will benefit from including additional known environmental and genetic factors. An important environmental factor is vocal occupation, i.e. the load which the vocal organ experiences on a day-to-day basis. Voice use represents a considerable occupational hazard (48) and could affect the organ’s shape via remodeling.

### Functional consequences of sex-specific morphotypes

Separate cluster analyses among females and among males identified different patterns of clustering. While in males the clusters differed by age, in females they differed by age and BMI. Again, it is notable that size of the cartilage is not an important factor explaining shape variation within sexes. 3D images were digitally morphed into cluster average images using a computer-automated algorithm. The approach appears to illustrate different laryngeal morphotypes, i.e. groups of individuals share large-scale shape features. The existence of laryngeal morphotypes among humans has been hypothesized, mostly in the context of structural adaptations for specific voice categories (6, 47, 49, 50). The sources of such morphological variation were largely unknown but this study demonstrated a quantitative approach validate cluster identity. It remains to be seen if cluster models can be trained so that new subjects can be assigned a certain morphotype.

The observed shape changes of the thyroid cartilage likely contribute to functional differences, for example vocal changes. To produce a voice, vocal folds are set into self-sustained airflow-induced vibrations (24). The vocal folds are housed inside the larynx and serve as sound source. Vocal fold vibrations not only depend on composition and viscoelastic properties but also vocal fold shape, position and glottal opening (24, 51, 52). Shape, position and glottal opening are critically dependent on thyroid cartilage shape (53).

The current study informs also the debate of the role of vocal organ size in determining voice features. Voice fundamental frequency is sometimes simplistically described to be a function of larynx size but should take vocal fold characteristics into account (37). The size of the cartilaginous framework can be misleading. Thyroid cartilage shape features (e.g., relative antero-posterior extension and angulation between the thyroid laminae) that affect the shape of the vocal folds, and therefore fundamental frequency, are size-independent within sexes. Although both sexes show size differences in the larynx related to selection for larger body size, selection on males for larger larynx size related to the production of lower fundamental frequency voices (54) does not result in similar changes in the female larynx. In other words, the difference we see in the shape between sexes has evolved in the two sexes through sexual selection rather than representing an extension of a broader size-dependent scaling of shape. Both the larger size and relatively longer antero-posterior shape of the cartilage contribute to longer vocal folds documented in males compared to female (13) which contributes to the typically deeper voice in males. However, vocal fold shape, thickness and composition contribute as well. Moreover, the taller and narrower thyroid cartilage in males contributes to a longer and narrower epilaryngeal tube with consequences on vocal tract filtering (55). Both sound source and vocal tract filter are also critical for evaluating characteristic voice disorders, or typical age-related vocal changes (56–59).

Shape changes of the thyroid cartilage can also affect the breathing function of the larynx. The larynx has been implicated in the etiology of obstructive sleep apnea (OSA) (60, 61), a sleep disorder that is caused by the recurring collapse of soft tissue structures in the upper airway. The underlying causal mechanism is related to motor and/or sensory dysfunction, and among the most important predisposing factors are obesity, male sex and age (62). While thyroid cartilage shape has been long known to reflect sex differences, this is the first study to provide statistical support for a role of age and BMI on larynx shape. Age was implicated in shape morphotypes within males and the BMI is a source of laryngeal shape variation in females. Indeed, males (relative to females), women with higher BMI and older males all have relatively more acute angles between the left and right thyroid laminae thereby narrowing the airway (Figure 4). Shape of the cartilaginous framework is likely associated with remodeling of the soft tissue inside and above the larynx. For example, a change in the alpha angle may lead to remodeling of the soft tissue in and above the larynx with consequences for airway patency and glottal sufficiency.

### The lifelong growth of the male larynx

We found that the overall size of the male thyroid cartilage increased at a rate of about 0.7% per decade. Centroid size of the female thyroid cartilage did not increase in a systematic age-dependent manner. As far as we know, a post-pubertal size increase of the larynx has not been reported in previous studies. In general, all soft and hard tissues of the human head and neck region continue to change in shape or size after puberty (63, 64). However, two other cranial cartilaginous structures increase in size throughout life similar to the male larynx. The human outer ear demonstrates sexual size differences and a lifelong size increase in both sexes (65–67). Likewise, the external nose is also sexually dimorphic and its dimensions increase in both sexes up to an age of at least 65 years (68). Testosterone sensitivity is the main driver of the laryngeal growth spurt during puberty. It remains to be seen whether an individual’s hormone profile continues to affect larynx shape (Abitol et al. 1999) or whether other mechanisms are responsible for the post-puberty continued growth and shape changes.

### Post-puberty trajectories of shape change

Sexual dimorphism of the larynx develops during adolescence between 12 and 18 years of age (13). Maturational growth of the human body ends between 18 and 25 years of age (69). Results from the current study suggest that laryngeal shape and size continue to change beyond the adolescent period. Those changes are different in females and males, and various environmental factors might be responsible. The post-pubertal trajectory of the male larynx reflects shape changes related to scaling. In contrast, post-pubertal shape changes of the female larynx seem more complex. Multiple factors play a role, with several potential trajectories.

In general, shape and size changes of the human body may manifest through spurts of remodeling (70). The laryngeal shape change in adolescent males during puberty, for example, occurs relatively fast (13) and can be associated with notable voice problems (i.e., breaking voice) for extended periods of times before the adult normal male voice has manifested (71). Whether the observed shape and size changes between 18 and 88 years of age occur slowly or quickly cannot be deduced from the current approach.

A second period of hormonal changes is associated with permanent cessation of ovarian function (menopause) and a reduction of testis function in females and males, respectively. Vocal changes associated with menopause have been reported (72) Here we noted that a prominent shape change was noted in the female Cluster 3 with the highest average age (Table 6, Figure 5B). Future work may investigate whether the shape changes of laryngeal cartilages are caused by hormonal changes, how fast they may manifest, how they relate to soft tissue and airway remodeling and how they affect acoustic properties. Rodent models could provide valuable information (40).

### Conclusions

We identified multiple sources of laryngeal shape variation including sex, age and BMI. The sources act differently between the sexes and generate morphotypes, i.e. clusters of similar shape, within females and within males. Interestingly, the male larynx demonstrated a lifelong size increase. Growth trajectories play a central role in life course epidemiology. They can serve as markers of prenatal or childhood development and/or for determinants of adult health outcomes. Interestingly, both, age and BMI, are linked to numerous other anatomical variation and are identified risk factors for laryngeal pathologies (73).

The current approach of 3D reconstruction of individual laryngeal elements, shape analysis through geometric morphometrics and thin-plate spline warping to extract biomechanically relevant shape variables overcomes two important challenges of laryngeal shape analysis. First, the approach is no longer dependent on cadaver measurements but can collect data from very large samples of individuals and thereby control for important factors. Second, the approach is independent from anatomical landmarks and is able to deliver measurements that are relevant for biomechanical modeling purposes (27). This study takes an important first step of integrating 3D imaging and explores functionally important variables (Figure 4). Next, anatomical differences can be visualized and then studied through computational modeling (37, 38). The current approach promises practicability and potential data for data mining through computer-automated analysis and integration with finite element models to link shape variation with function since the analysis is relatively fast. Any new subject can be assigned to a morphotype in less than one hour after the CT scans have been taken.

## Methods

### Human subject sample

All methods were performed in accordance with the guidelines and regulations by Mayo Clinic and Midwestern University specified in a collaboration agreement. This retrospective study was approved by the Mayo Clinic Institutional Review Board, and the need for informed consent was waived. The radiology database was queried to identify adult patients (age ≥ 18 years) who had undergone consecutive neck CT scans for clinically-indicated purposes. The electronic medical records were reviewed for these potential study subjects, and individuals with a history of any of the following were specifically excluded (**1**) head and neck cancer, (**2**) irradiation of the head and neck region, (**3**) surgery involving the skull base, neck, or upper aerodigestive tract (with the exception of tonsillectomy), (**4**) laryngeal dysfunction documented on physical exam or symptoms suggesting a possible laryngeal disorder (e.g. hoarseness, aspiration), (**5**) treatment for a systemic malignancy within the past 12 months (due to potential confounding effect on weight).

To ensure broad sampling for the variables of interest, subjects were consecutively recruited in a stratified fashion based upon sex (male, female), age (18-28; 28-38; 38-48; 48-58; 58-68; 68-78; 78-88 years), and body mass index (BMI) (<25; 25-29.9; ≥30 kg/m^2^). The BMI, weight/height^2^, is correlated with body fat used by medical professionals to screen for under- or over weight status (74). To be eligible for inclusion, subjects had to have height and weight documented within 30 days of the neck CT acquisition. Five subjects were enrolled into each of these 42 category-combinations for a total of 210 subjects.

Body height is also an important determinant of larynx size (75), which is itself an important determinant of laryngeal function (37). Body height was not included in the stratification criteria for subject sampling because the projected sample size would have been very large. However, the post hoc examination of the sample revealed considerable variation in body height, which allowed the investigation of its effect on larynx shape and size. We included height as variable in the investigation of variation within females and males, but excluded height from the analysis of the mixed-sex sample given its strong covariation with sex (Pearson correlation, r=0.71, P<0.001).

#### Imaging, segmentation, 3D surface renderings, landmark coordinates and centroid size

Neck CT scans were reviewed by a board-certified neuroradiologist to ensure good image quality and technical adequacy. Although performed using different multidetector helical CT scanner models, the neck CT’s were all acquired in supine position using 120 kVp with an mAs that was adjusted automatically by the scanner to maintain adequate penetration in the setting of variable body habitus. As a minimum, reconstructed z-axis slice thickness had to be < 0.7 mm with an axial field of view ≤ 28 cm and a matrix size of 512. Images were rendered with a standard soft tissue kernel. Subjects were eligible for inclusion regardless of whether or not intravenous contrast was administered for the CT. All CT datasets were anonymized prior to export for shape analysis.

Reconstructed image stacks were imported into AVIZO software (v. Lite 9.0.1) and thyroid and cricoid cartilages were segmented manually. Segmentation refers to the process of generating surface renderings of anatomical structures. Derived 3D surfaces (stereolithography format) of all specimens are available on Morphobank (www.morphobank.com), Project 4345.

The 3D surfaces of both cartilages were then imported to the *geomorph* package in the statistical software program R 3.4.4 (R Core Team 2015) (76) to perform surface landmark placement. Landmarks were collected from surface renderings in the form of three-dimensional coordinates (77). The landmark and semilandmark protocol captured variation in homologous structures, including structural margins and surfaces such as the thyroid lamina. Our previous work showed that 100 surface semilandmarks are sufficient to capture shape variation in the thyroid cartilage to achieve comprehensive coverage (77).

Centroid size was used as a measure of cartilage size. Centroid size is computed as the square root of the sum of squared distances of each landmark from the centroid of the cartilage landmarks, whose location is obtained by averaging the *x, y* and *z* coordinates of all landmarks (78).

### Analysis

The first step in the geometric morphometric shape analysis was superimposition of the landmark configurations to remove differences in raw scale, orientation and position in the global coordinate system. This was accomplished via generalized Procrustes analysis (GPA) which extracts shape information from the raw coordinate data by translating landmark configurations to a common location, scaling configurations to unit centroid size, and rigidly rotating configurations to minimize squared distances among corresponding landmarks (79). As the resulting shape variables exist in a non-Euclidean shape space, we projected them to a linear tangent space prior to statistical analysis. This produces a set of transformed coordinates that reflect shape differences among cartilages independent of size.

A series of principal components analyses (PCA) were then conducted on the full sample, males only and females only to reduce the dimensionality of the data and summarize the main patterns of shape variation. We evaluated only those PCs that cumulatively explained 85% of the variation. Each PC excluded from the analysis explained <0.5% of variation. As a result, we used 22 (men and women combined), 24 (men) and 18 (women) PCs, respectively, to explore patterns of similarity.

We began by exploring the relationship among shape variables, age, BMI, centroid size and height using Pearson correlation analyses. We then used multiple analysis of variance (MANOVA) with superimposed shape coordinates as dependent variables and sex, age, BMI, and height (in sex specific analyses only) as main effects as well as their two-way interactions using the ProcD.lm function in the geomorph package for R. Type II sums-of-squares was applied to evaluate each main effect independent of the other main effects. Statistical significance is based on randomization of residuals via permutation, an appropriate approach for high-dimensional data such as landmarks and semi landmarks (76).

K-means clustering (R Core Team 2021) was performed on the limited subset of principal components described above to assess whether cartilages formed distinct clusters in the shape space. The clustering method is designed to create homogeneous groups of individuals (e.g., clusters) that are maximally different from other clusters. The silhouette coefficient was used to estimate the number of relevant clusters (80) using the cluster package in R (81). Its calculation provides a measure of how similar an object is to its own cluster (cohesion) compared to other clusters (separation). The Silhouette Score was calculated as: (b-a)/max(a,b), where a is the average intra-cluster distance (average distance between each point within a cluster) and b is the average inter-cluster distance (the average distance between all cluster centroids). The Silhouette Score can range between −1 and 1 where a high value indicates the object is well matched to its own cluster and poorly matched to neighboring clusters. Values near zero indicate that the distance between two clusters is not significant. Based on silhouette scores, the mix-sex sample was analyzed with two clusters; three clusters were indicated for the male sample and four clusters for females (Supplemental Figure S1). We compared the average values of the age, BMI or body height variables within each cluster using t-tests or ANOVA to assess whether the observed clustering of shape axes corresponds to variation in these factors, as appropriate.

### Warping

To illustrate what anatomical information was encoded by individual shape axes, by regression models or by individual clusters in the cluster analyses, a thin-plate spline method was used to warp the thyroid cartilage into an estimated mean shape for a set of aligned specimens. This was achieved with geomorph’s *warpRefMesh* function (76). The function uses the thin-plate spline method to warp a surface model of the mean shape for a set of aligned specimens, into the shape defined by a second set of landmark coordinates. For example, the latter coordinates may represent the mean of a subsample, such as males, or may be the scaled coefficients from the regression of shape on age added to the mean configuration to represent the effect of aging.

A challenge is the translation of statistical shape differences into mechanically relevant information. We extracted two angular and five linear shape measurements with biomechanical relevance for vocal organ function from warped surface reconstructions to more directly investigate the biomechanical consequence of shape differences documented in the current study.

## Author contributions

Conceptualization: TR, JH

Methodology: TR, JH, KB, AS

Investigation: TR, KB, AS Visualization: TR, KB Supervision: TR

Writing—original draft, review & editing: TR, JH, KB, AS

## Data and materials availability

Anonymized DICOM data for the neck CT scans cannot be made publicly available because of the potential to create a three-dimensional rendering of the face, which is considered to be sensitive patient health information that must be protected. Derived 3D surface reconstructions (stereolithography format) of all thyroid cartilage specimens are available on Morphobank, Project 4345.

---

**Supplemental Figure S1:**
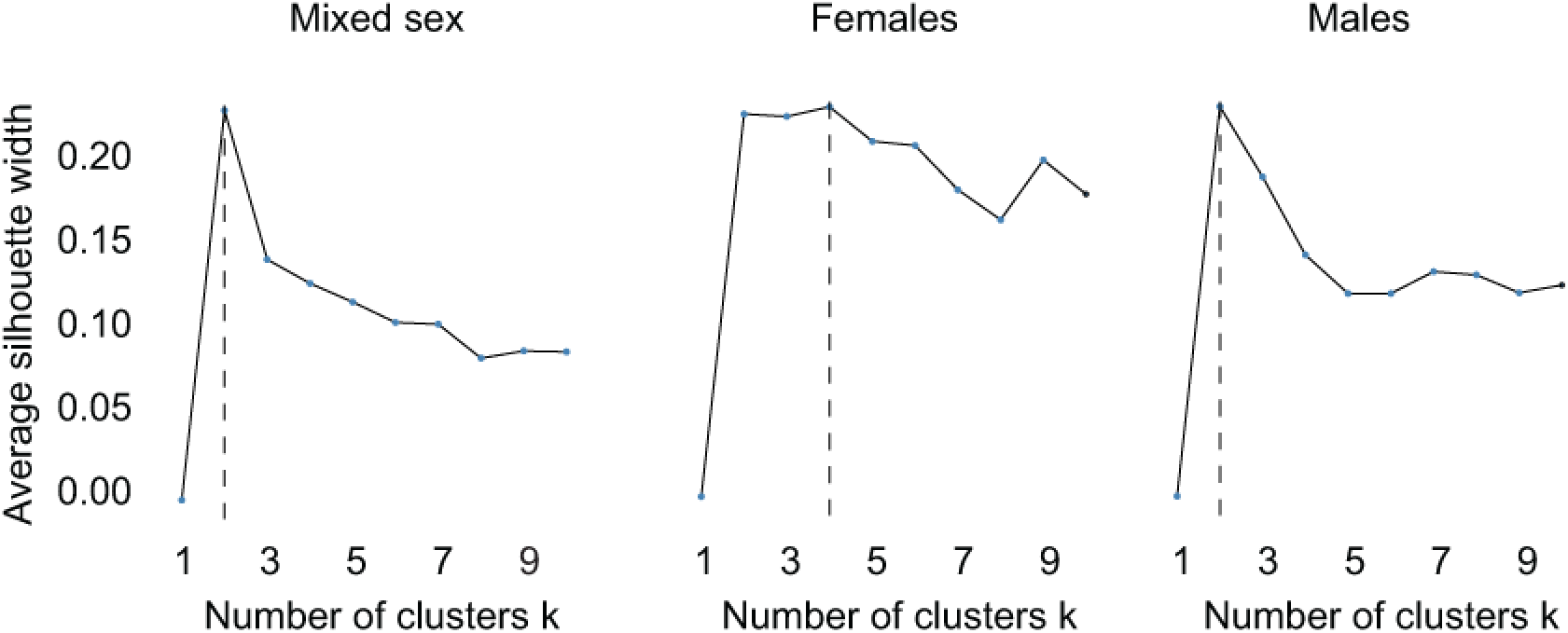
Optimal number of clusters. The silhouette score for males and females combined, recommends 2 clusters. The silhouette analysis (silhouette score) recommends 4 clusters for females and 2 to 3 clusters for males.

